## Supplementary materials for "*Bacillus velezensis* HBXN2020 alleviates *Salmonella* Typhimurium infection in mice by improving intestinal barrier integrity and reducing inflammation"

**Supplementary Figures and legends**

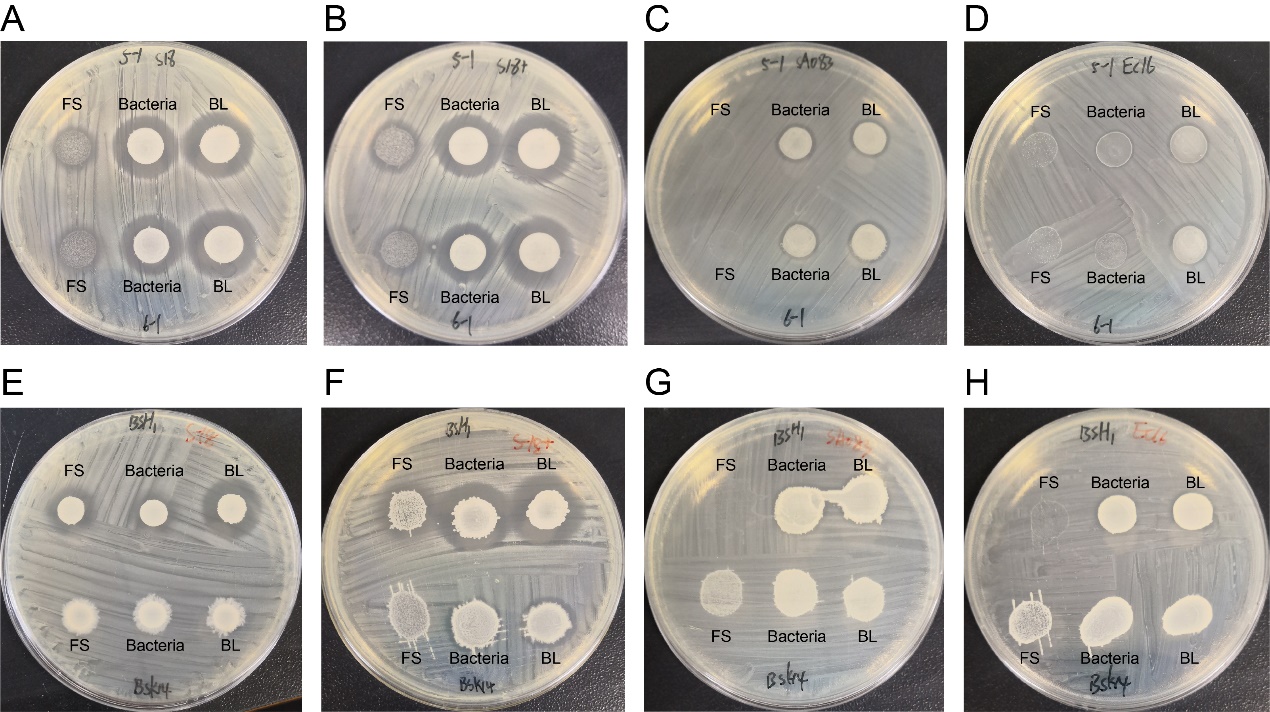

**Supplemental Figure 1. Spot-on-plate assay of *Bacillus***. (**A**) and (**E**) indicator bacteria are *S. aureus* S18; (**B**) and (**F**) indicator bacteria are *S. aureus* S18+; (**C**) and (**G**) indicator bacteria are *Salmonella* SA083 (*S*. Enteritidis SE006); (**D**) and (**H**) indicator bacteria are *E. coli* EC16 (*E. coli* EC016). FS：Fermentation supernatant；BL：Bacteria liquid

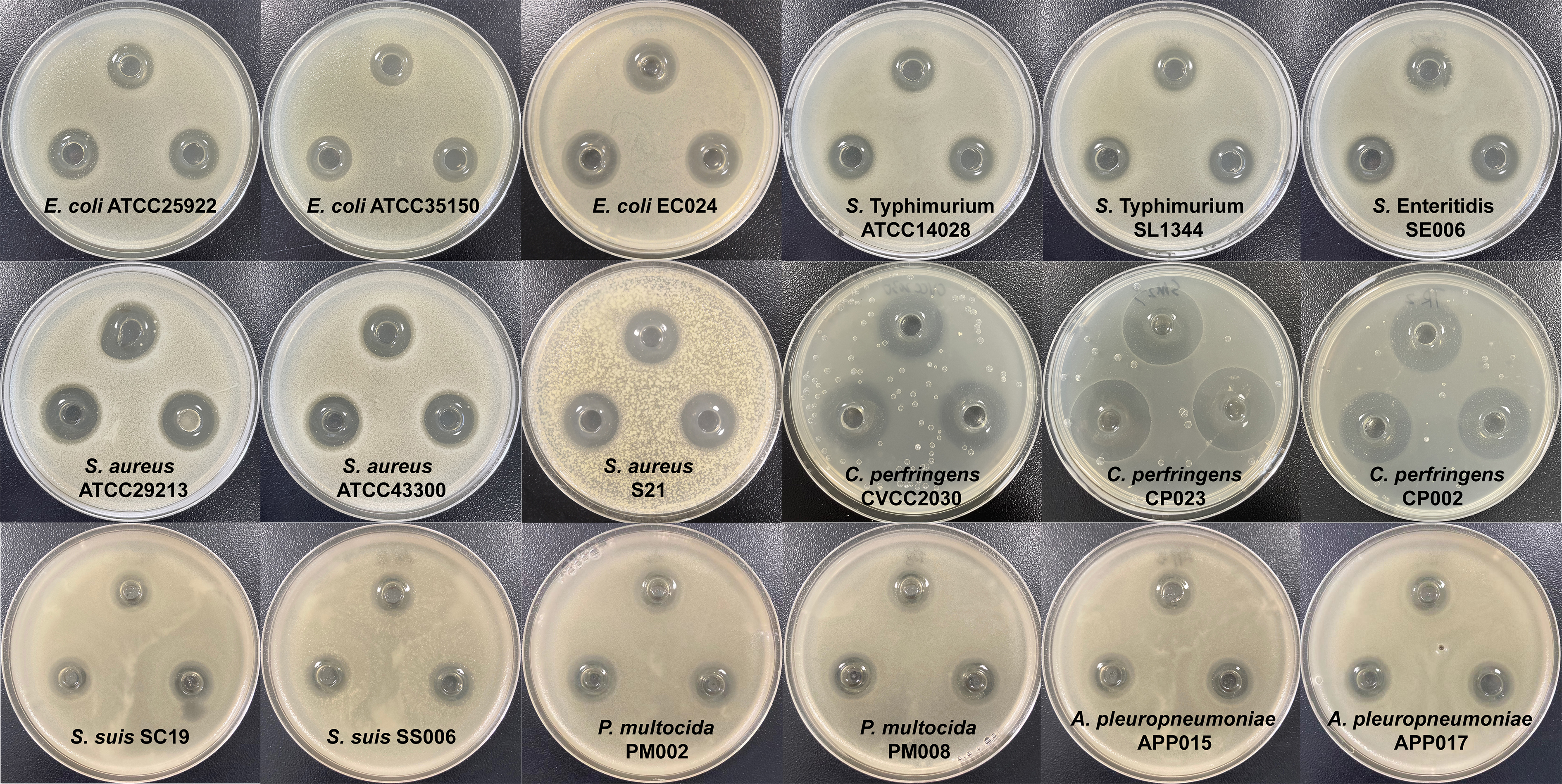

**Supplemental Figure 2. *In vitro* antibacterial test of *B. velezensis* HBXN2020**. The in vitro antagonistic activity of *B. velezensis* HBXN2020 against 18 indicator strains (pathogens) was tested using agar well-diffusion method. The fermentation supernatant of *B. velezensis* HBXN2020 was filtered with a 0.22 μm filter and add it to different Oxford cups. All plates were cultured at 37°C for 16 h before observing the inhibition zone, and the diameter of the inhibition zone were measured by vernier caliper. The clear zone was expressed the antagonistic activity.

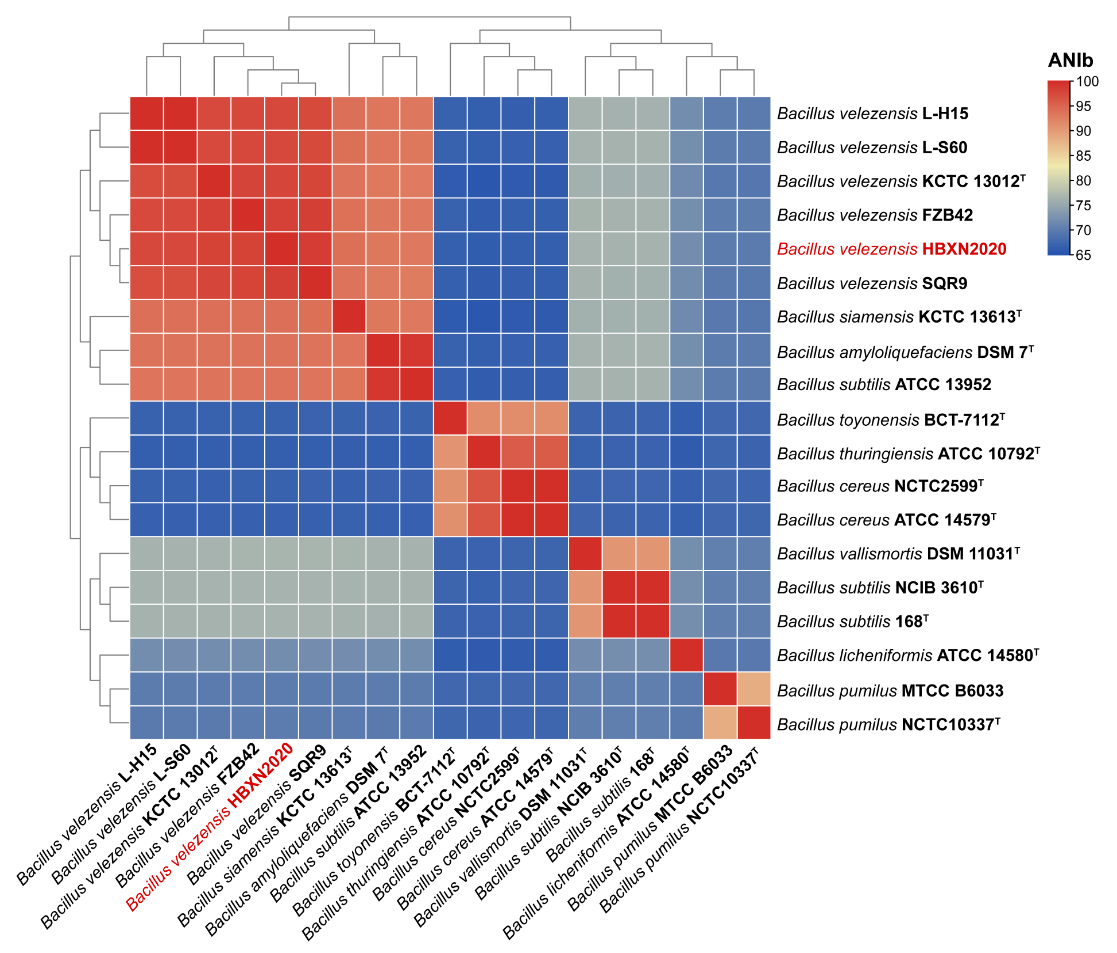

**Supplemental Figure 3. *B. velezensis* HBXN2020 genome ANI analysis**. The heatmap based on the ANIb value of strain HBXN2020 and other *Bacillus* species. The *B. velezensis* HBXN2020 was labeled in red letters.

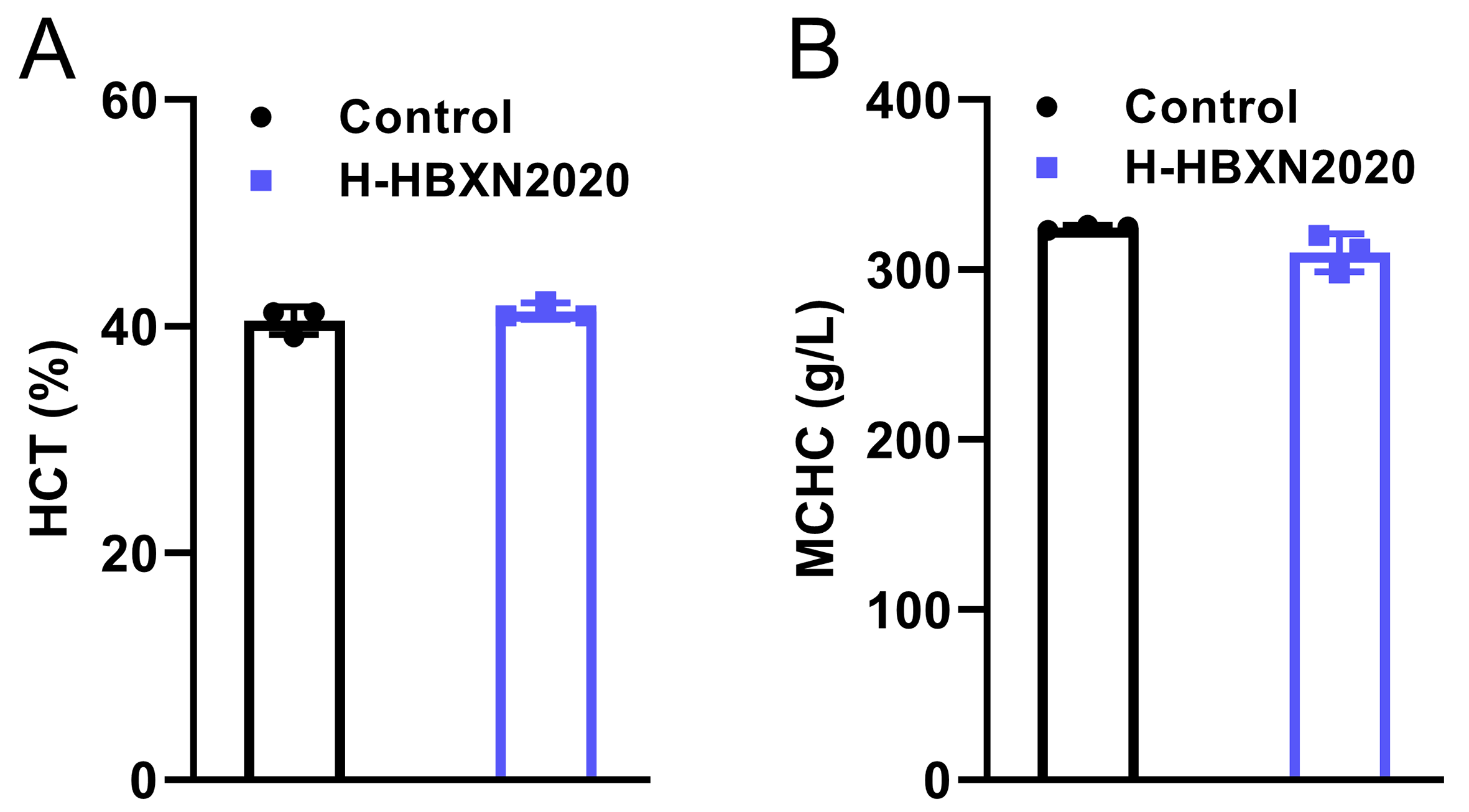

**Supplemental Figure 4. Hematological parameters**. Blood routine parameters of mice in the control group and high dose *B. velezensis* HBXN2020 group. (**A**) Hematocrit (HCT), (**B**) Mean corpuscular hemoglobin concentration (MCHC).

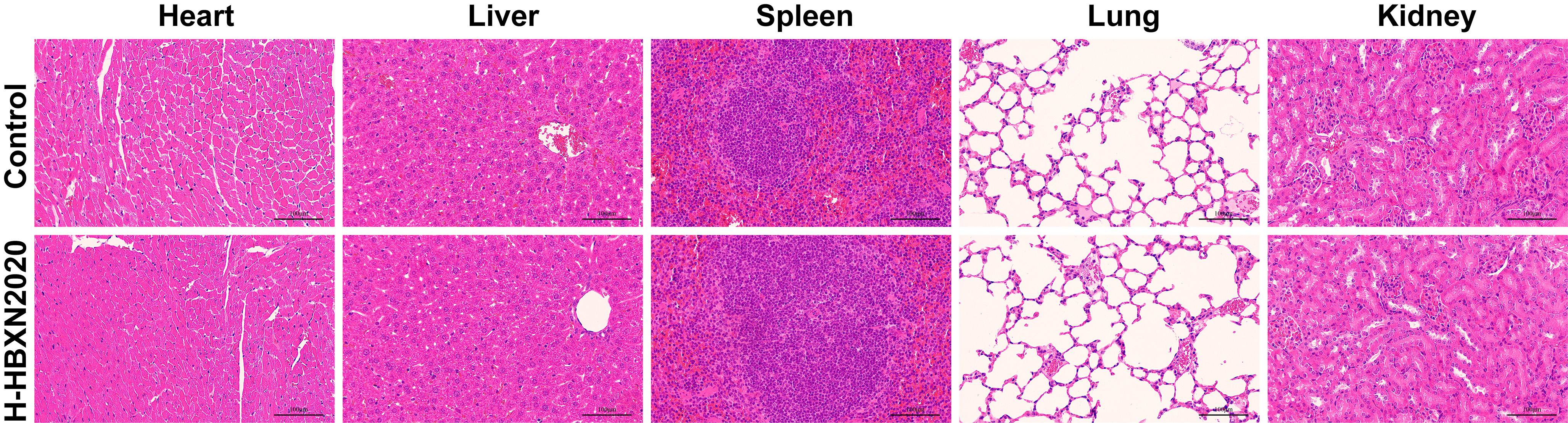

**Supplemental Figure 5. H&E staining of the intestine, heart, liver, spleen, lung and kidney**. Tissues were obtained from mice in the high dose *B. velezensis* HBXN2020 group at 15 days. PBS was used as a control. Scale bars: 100 µm.

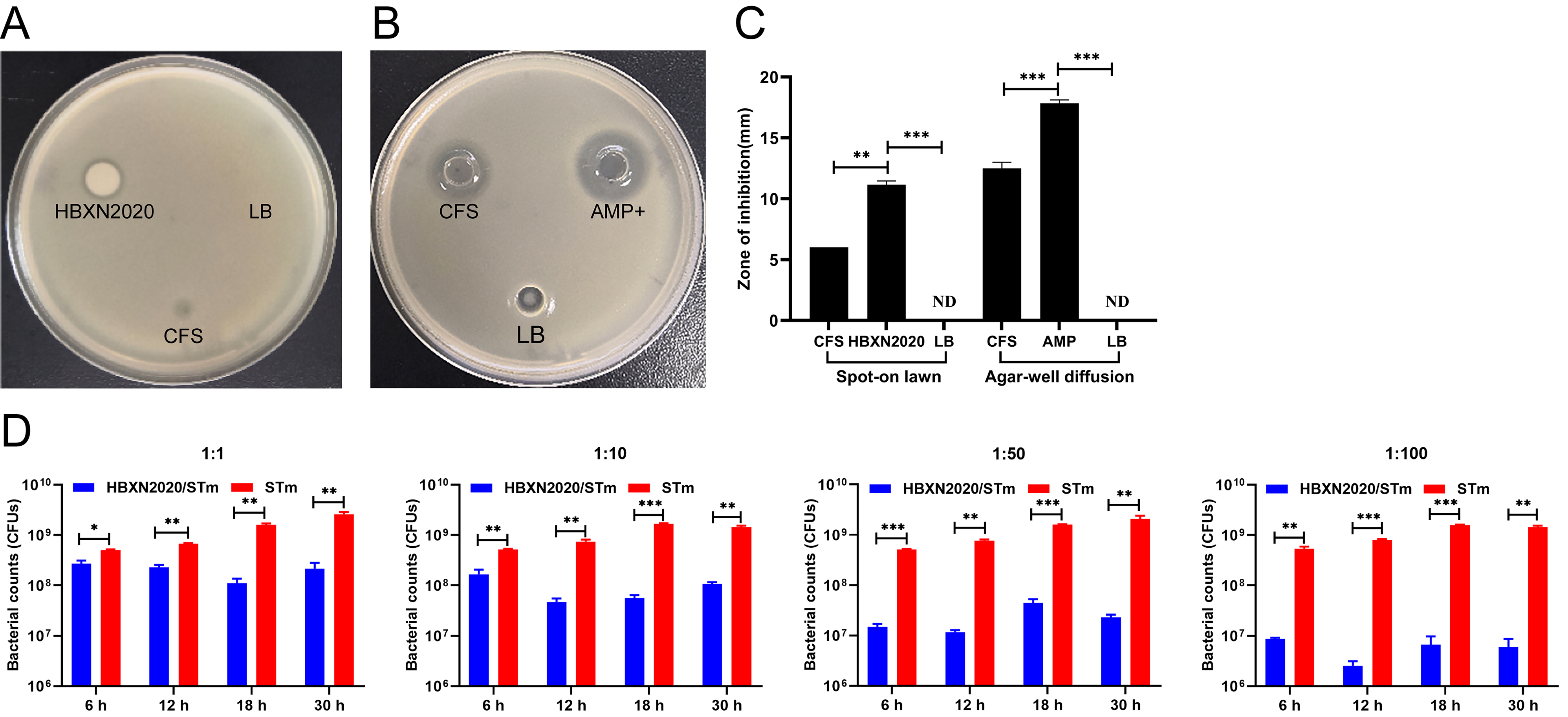

**Supplemental Figure 6. In vitro antagonistic activity of *B. velezensis* HBXN2020 against *S*. Typhimurium ATCC14028 in liquid and solid media.** The antagonistic *Salmonella* activities of HBXN2020 cell-free supernatant (CFS) were determined using spot-on lawn assay (**A**) and agar well diffusion assay (**B**). (**C**) Diameter of inhibition zone in the spot-on lawn assay and agar well diffusion assay. Data were shown as mean values ± SEM (n = 3). Each group was repeated three times. AMP+, ampicillin as a positive control, LB medium as a negative control. ND, no detectable. (**D**) *S*. Typhimurium ATCC14028 (STm) were co-incubated with *B. velezensis* HBXN2020 in LB medium at various ratios at 37°C. The survivals of STm were examined at indicated time points by bacterial counting on selective agar plates. Data were shown as mean values ± SEM (n = 3). Statistical signifcance was evaluated using Student’s t-test. *, *p* < 0.05, **, *p* < 0.01, and ***, *p* < 0.001.

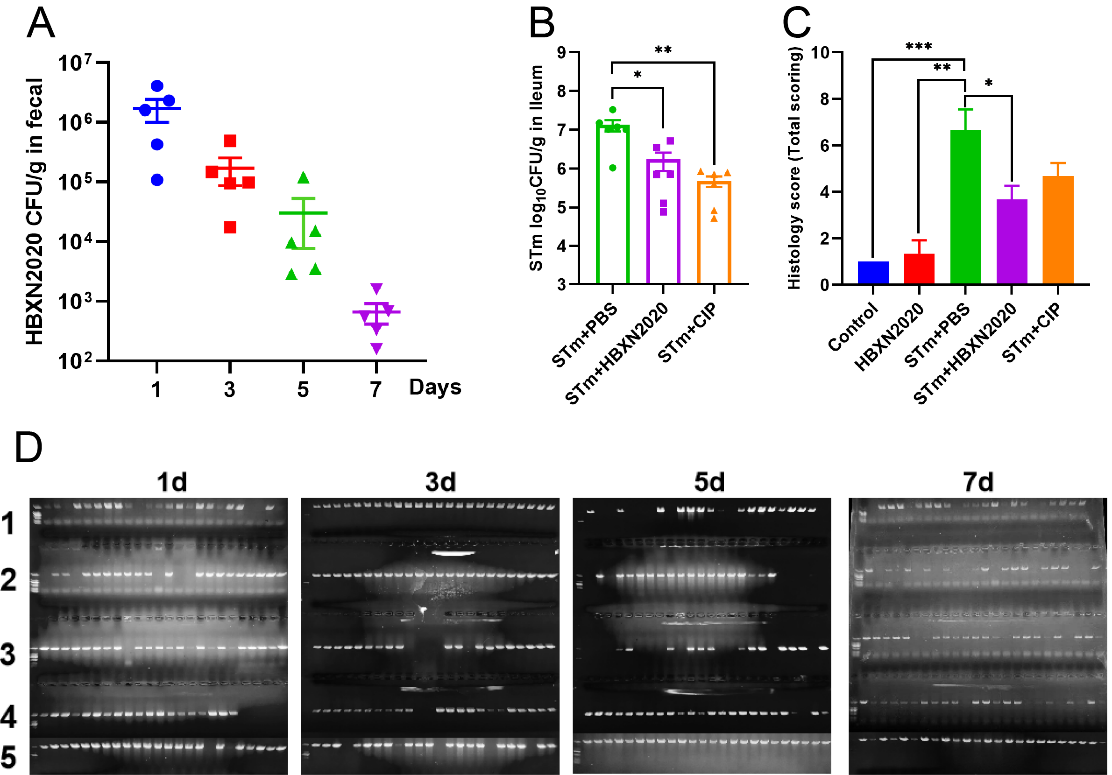

**Supplemental Figure 7. The effect of therapeutic *B. velezensis* HBXN2020 on the bacterial load in feces of mice and the histological score of the colons**. (**A**) The bacterial load of *B. velezensis* HBXN2020 in mouse feces. The quantity of *B. velezensis* HBXN2020 was identified by combining bacterial count and PCR (n = 5). Data were shown as mean values ± SEM. (**B**) The bacterial loads of *S*. Typhimurium ATCC14028 in ileum. The ileum was harvested and then homogenized. One hundred microliters of each sample performed a serial of 10-fold dilutions and spread on selective agar plates and incubated at 37°C for 12 h before bacterial counting. (**C**) Histological scores of colons (n = 3). (**D**) *B. velezensis* HBXN2020 single colony PCR identification results. Statistical signifcance was evaluated using one-way analysis of variance (ANOVA) with Tukey’s multiple comparisons test (*, *p* < 0.05, **, *p* < 0.01, and ***, *p* < 0.001).

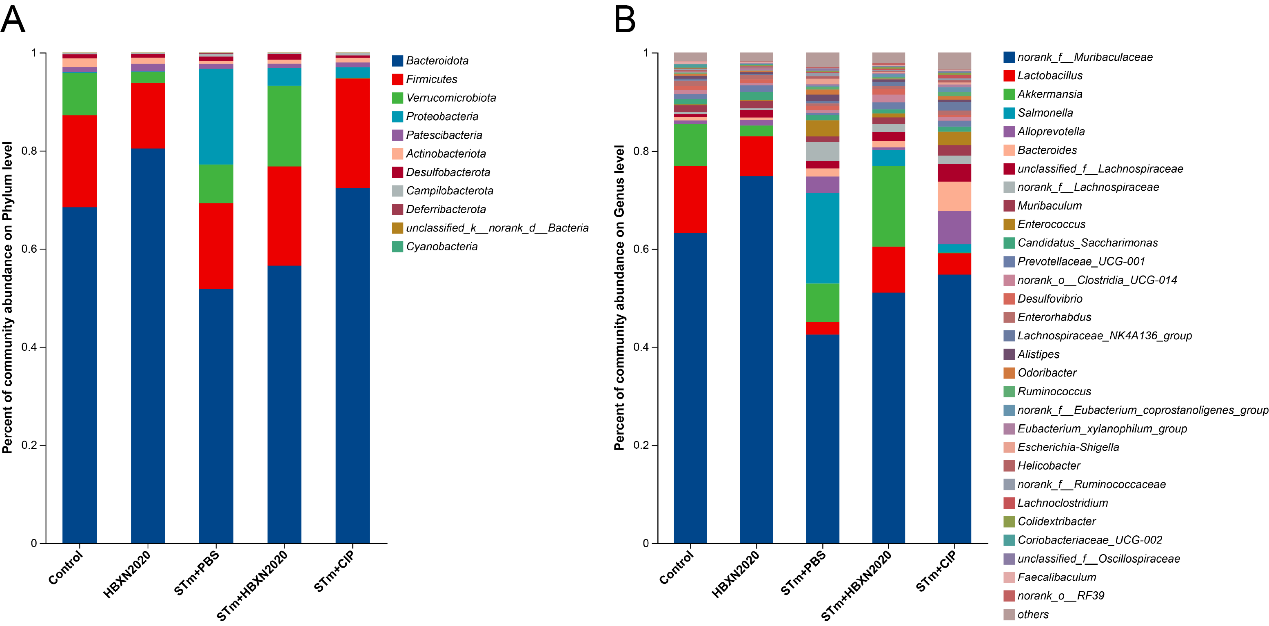

**Supplemental Figure 8. Analysis of the community compositions of colonic microbiota**. (**A**) Community compositions analysis at the phylum level. This bar graph shows the 15 most abundant communities. (**B**) Community compositions analysis at the genus level. This bar graph shows the communities of the top 30 genera.

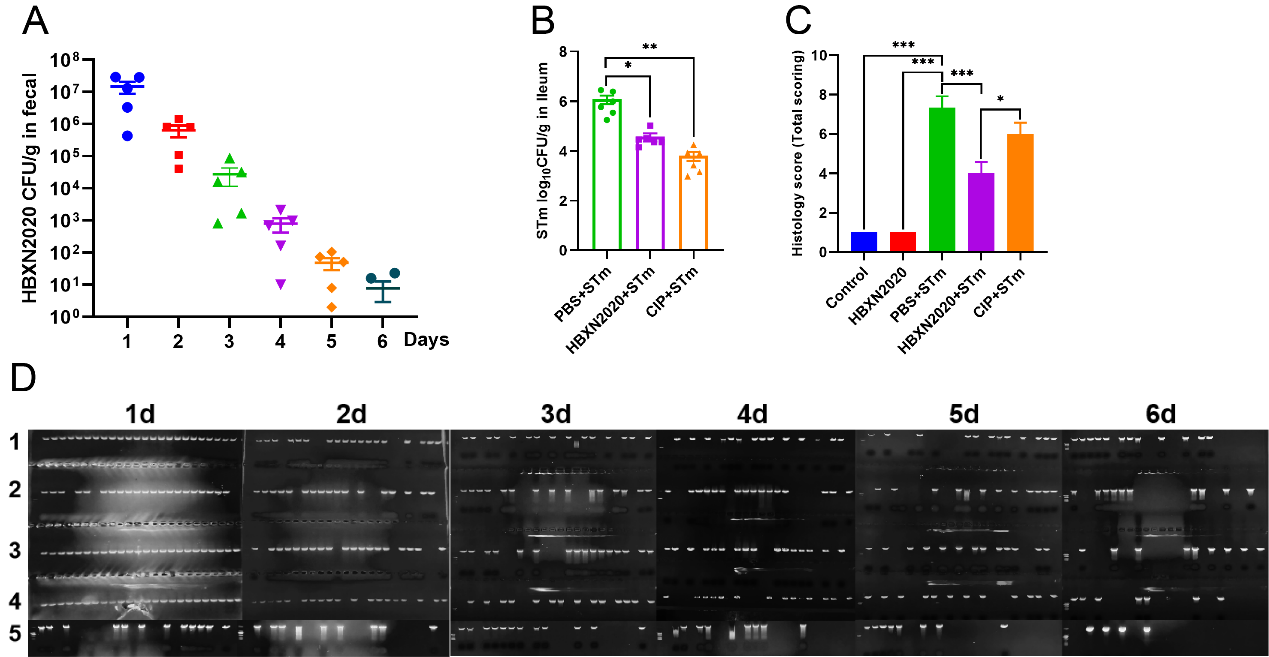

**Supplemental Figure 9. The effect of prophylactic *B. velezensis* HBXN2020 on the bacterial load in feces or ileum of mice and the histological score of the colons.** (**A**) The bacterial load of *B. velezensis* HBXN2020 in mouse feces. The quantity of *B. velezensis* HBXN2020 was identified by combining bacterial count and PCR (n = 5). Data were shown as mean values ± SEM. (**B**) The bacterial loads of *S*. Typhimurium ATCC14028 in ileum. The ileum was harvested and then homogenized. One hundred microliters of each sample performed a serial of 10-fold dilutions and spread on selective agar plates and incubated at 37°C for 12 h before bacterial counting. Statistical signifcance was evaluated using Student’s t-test (*, *p* < 0.05, **, *p* < 0.01, and ***, *p* < 0.001). (**C**) Histological scores of colons (n = 3). (**D**) *B. velezensis* HBXN2020 single colony PCR identification results. Statistical signifcance was evaluated using one-way analysis of variance (ANOVA) with Tukey’s multiple comparisons test (*, *p* < 0.05, **, *p* < 0.01, and ***, *p* < 0.001).

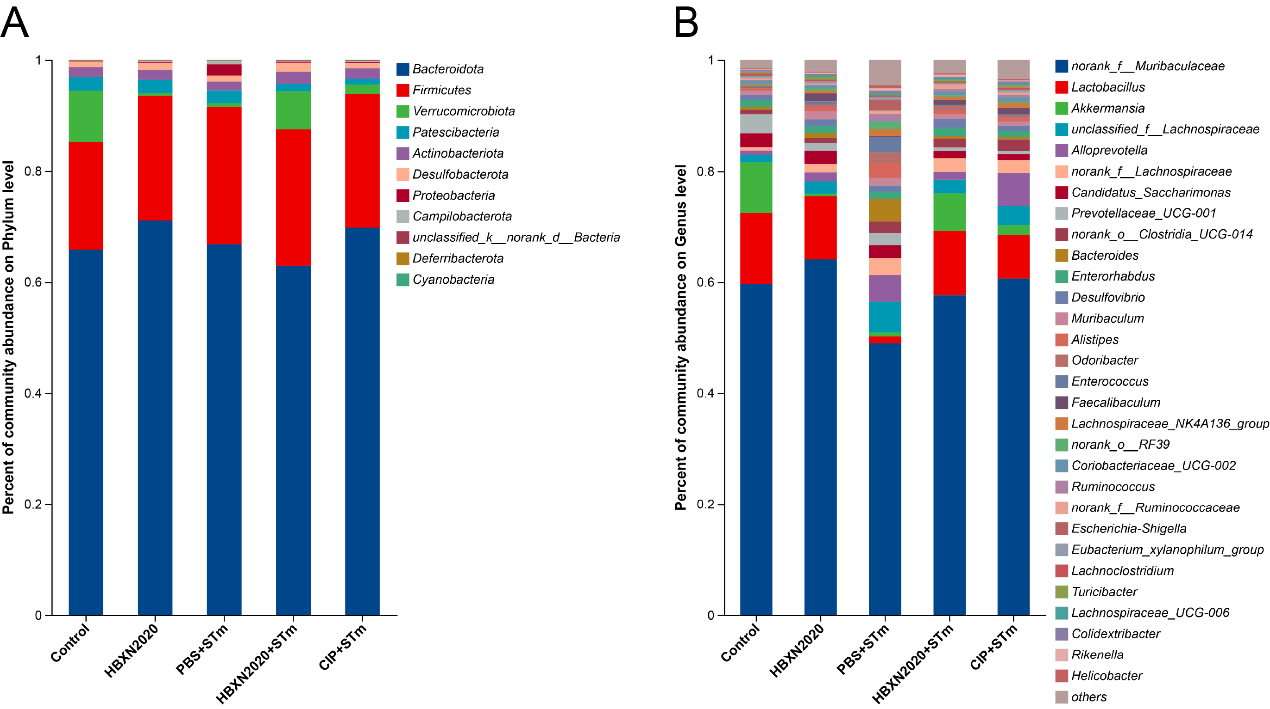

**Supplemental Figure 10. Analysis of the community compositions of colonic microbiota**. (**A**) Community compositions analysis at the phylum level. This bar graph shows the 15 most abundant communities. (**B**) Community compositions analysis at the genus level. This bar graph shows the communities of the top 30 genera.

**Supplementary Tables**

**Supplementary Table 1. Determination of antibacterial activity of different *Bacillus***

| **Bacteria**  **and serovars** | **Strain no.** | B. ***subtilis***   **BSH1** | ***B. velezensis***  **HBXN2020** | ***B. amyloliquefaciens***  **6-1** | B. ***licheniformis***   **BSK14** |
| --- | --- | --- | --- | --- | --- |
| *E. coli* | EC006 | - | ++ | + | - |
|  | EC016 | - | +++ | + | - |
|  | EC022 | - | ++ | - | - |
|  | EC024 | - | +++ | ++ | - |
|  | ATCC  35150 | - | +++ | ++ | - |
|  | ATCC  25922 | - | +++ | ++ | - |
| *S.* Typhimurium | ST001 | - | +++ | ++ | - |
|  | ST002 | - | ++ | +++ | - |
|  | ST003 | - | + | + | - |
|  | ST004 | - | +++ | + | - |
|  | ST005 | - | +++ | ++ | - |
|  | ST006 | - | +++ | +++ | - |
|  | ST007 | - | +++ | ++ | - |
|  | SL1344 | - | +++ | ++ | - |
|  | ATCC  14028 | - | +++ | + | - |
| S. Enteritidis | SE001 | - | +++ | ++ | - |
|  | SE002 | - | +++ | ++ | - |
|  | SE003 | - | +++ | ++ | - |
|  | SE004 | - | +++ | +++ | - |
|  | SE005 | - | ++ | ++ | - |
|  | SE006 | - | +++ | ++ | - |
| *S. aureus* | S1 | ++ | +++ | +++ | +++ |
|  | S2 | ++ | +++ | +++ | +++ |
|  | S3 | ++ | +++ | +++ | ++ |
|  | S4 | ++ | +++ | +++ | ++ |
|  | S5 | ++ | +++ | +++ | +++ |
|  | S6 | +++ | +++ | +++ | +++ |
|  | S10 | ++ | ++ | +++ | +++ |
|  | S11 | ++ | +++ | +++ | ++ |
|  | S12 | ++ | ++ | +++ | ++ |
|  | S13 | ++ | ++ | + | +++ |
|  | S14 | ++ | ++ | ++ | +++ |
|  | S15 | +++ | +++ | +++ | ++ |
|  | S16 | +++ | ++ | ++ | ++ |
|  | S17 | +++ | ++ | +++ | ++ |
|  | S18 | ++ | +++ | +++ | +++ |
|  | S19 | ++ | ++++ | +++ | +++ |
|  | S20 | ++ | ++ | +++ | ++ |
|  | S21 | ++ | +++ | ++ | +++ |
|  | ATCC  43300 | ++ | +++ | +++ | +++ |
|  | ATCC  25923 | ++ | +++ | +++ | ++ |
|  | ATCC  29213 | +++ | +++ | +++ | ++ |
| *S. suis* | SS6 | ++ | +++ | +++ | + |
|  | SS12 | ++ | +++ | +++ | ++ |
|  | SS54 | +++ | ++ | ++ | ++ |
|  | SS55 | ++ | ++ | ++ | ++ |
|  | SS57 | ++ | ++ | ++ | ++ |
|  | SS58 | +++ | +++ | +++ | ++ |
|  | SS59 | ++ | +++ | ++ | ++ |
|  | SS60 | +++ | ++ | +++ | +++ |
|  | SS61 | ++ | +++ | +++ | ++ |
|  | SS62 | ++ | ++ | ++ | ++ |
|  | SS63 | ++ | ++ | ++ | +++ |
|  | SS64 | ++ | ++ | ++ | ++ |
|  | SC19 | ++ | ++ | +++ | ++ |
| *C. perfringens* | CP001 | ++ | +++ | +++ | +++ |
|  | CP002 | +++ | ++++ | +++ | +++ |
|  | CP003 | ++ | +++ | +++ | +++ |
|  | CP004 | ++ | +++ | +++ | +++ |
|  | CP005 | ++ | +++ | +++ | +++ |
|  | CP006 | +++ | +++ | +++ | +++ |
|  | CP007 | ++ | +++ | +++ | +++ |
|  | CP008 | ++ | +++ | +++ | +++ |
|  | CP009 | ++ | +++ | +++ | +++ |
|  | CP010 | ++ | +++ | +++ | +++ |
|  | CP011 | ++ | +++ | +++ | +++ |
|  | CP012 | ++ | +++ | +++ | +++ |
|  | CP013 | ++ | +++ | +++ | +++ |
|  | CP014 | ++ | +++ | +++ | +++ |
|  | CP015 | ++ | +++ | +++ | +++ |
|  | CP016 | ++ | +++ | +++ | +++ |
|  | CP017 | ++ | +++ | +++ | +++ |
|  | CP018 | ++ | +++ | +++ | +++ |
|  | CP023 | ++ | ++++ | +++ | +++ |
|  | CVCC  2030 | ++ | ++++ | +++ | ++++ |
| *A. pleuropneumoniae* | APP015 | - | ++ | + | - |
|  | APP016 | - | ++ | ++ | - |
|  | APP017 | - | ++ | ++ | - |
|  | APP018 | - | ++ | + | - |
| *P. multocida* | PM002 | - | +++ | ++ | - |
|  | PM008 | - | +++ | +++ | - |

Note: -, No antibacterial activity; +, 0 < bacteriostatic diameter ≤ 5; ++, 5 < bacteriostatic diameter ≤ 15; +++, 15 < bacteriostatic diameter ≤ 20；++++, 20 < bacteriostatic diameter.

**Supplementary Table 2. HBXN2020 genome features**.

| Features | Value |
| --- | --- |
| Genome size (bp) | 3,929,792 |
| G+C content% | 46.5 |
| Protein coding genes | 3,744 |
| Gene average length (bp) | 928.06 |
| Plasmid | 0 |
| rRNA | 27 |
| tRNA | 86 |

**Supplementary Table 3. clusters of secondary metabolic synthesis genes in HBXN2020**

| BGC | BGC type | Length (bp) | BGC content (% Similarity) |
| --- | --- | --- | --- |
| 1 | NAD(P)/FAD-dependent oxidoreductase | 63978 | Surfactin (82%) |
| 2 | PKS-like | 41254 | Butirosin A / Butirosin B (7%) |
| 3 | Hypothetical protein | 17409 | Unknown |
| 4 | Lanthipeptide | 28889 | Unknown |
| 5 | 1-phosphofructokinase | 87836 | Macrolactin H (100%) |
| 6 | Competence/damage-inducible protein A | 109575 | Bacillaene (100%) |
| 7 | Zinc-binding alcohol dehydrogenase family protein | 134311 | Fengycin (100%) |
| 8 | LysM peptidoglycan-binding domain-containing protein | 19553 | Unknown |
| 9 | Terpene | 21884 | Unknown |
| 10 | T3PKS | 41101 | Unknown |
| 11 | TransAT-PKS-like | 106183 | Difficidin (100%) |
| 12 | NRPS | 51792 | Bacillibactin (100%) |
| 13 | Other | 41419 | Bacilysin (100%) |

**Supplementary Table 4. The bacterial strains used in this study.**

| **Strains** | **Strain ID number** | **Source** |
| --- | --- | --- |
| *Bacillus velezensis* (*B.* velezensis, NCBI no. CP119399) | HBXN2020 | Lab stock |
| *Escherischia coli* (*E. coli*) | ATCC25922 | ATCC |
|  | ATCC35150 | ATCC |
|  | EC024 | Lab stock |
|  | EC022 | Lab stock |
|  | EC016 | Lab stock |
|  | EC006 | Lab stock |
| *Salmonella enterica* serovar Typhimurium (*S.* Typhimurium, carry pET28a (+), kanamycin resistance) | ATCC14028 | ATCC |
|  | SL1344 | Lab stock |
|  | ST001 | Lab stock |
|  | ST002 | Lab stock |
|  | ST003 | Lab stock |
|  | ST004 | Lab stock |
|  | ST005 | Lab stock |
|  | ST006 | Lab stock |
|  | ST007 | Lab stock |
| *Salmonella enterica* serovar Enteritidis (*S.* Enteritidis) | SE006 | Lab stock |
|  | SE001 | Lab stock |
|  | SE002 | Lab stock |
|  | SE003 | Lab stock |
|  | SE004 | Lab stock |
|  | SE005 | Lab stock |
| *Staphylococcus aureus* (*S. aureus*) | ATCC29213 | ATCC |
|  | ATCC43300 | ATCC |
|  | ATCC25923 | ATCC |
|  | S21 | Lab stock |
|  | S1 | Lab stock |
|  | S2 | Lab stock |
|  | S3 | Lab stock |
|  | S4 | Lab stock |
|  | S5 | Lab stock |
|  | S6 | Lab stock |
|  | S10 | Lab stock |
|  | S11 | Lab stock |
|  | S12 | Lab stock |
|  | S13 | Lab stock |
|  | S14 | Lab stock |
|  | S15 | Lab stock |
|  | S16 | Lab stock |
|  | S17 | Lab stock |
|  | S18 | Lab stock |
|  | S19 | Lab stock |
|  | S20 | Lab stock |
| *Clostridium perfringens* (*C. perfringens*) | CVCC2030 | Lab stock |
|  | CP023 | Lab stock |
|  | CP001 | Lab stock |
|  | CP002 | Lab stock |
|  | CP003 | Lab stock |
|  | CP004 | Lab stock |
|  | CP005 | Lab stock |
|  | CP006 | Lab stock |
|  | CP007 | Lab stock |
|  | CP008 | Lab stock |
|  | CP009 | Lab stock |
|  | CP010 | Lab stock |
|  | CP011 | Lab stock |
|  | CP012 | Lab stock |
|  | CP013 | Lab stock |
|  | CP014 | Lab stock |
|  | CP015 | Lab stock |
|  | CP016 | Lab stock |
|  | CP017 | Lab stock |
|  | CP018 | Lab stock |
| *Streptococcus suis* (*S. suis*) | SC19 | Lab stock |
|  | SS006 | Lab stock |
|  | SS6 | Lab stock |
|  | SS12 | Lab stock |
|  | SS54 | Lab stock |
|  | SS55 | Lab stock |
|  | SS57 | Lab stock |
|  | SS58 | Lab stock |
|  | SS59 | Lab stock |
|  | SS60 | Lab stock |
|  | SS61 | Lab stock |
|  | SS62 | Lab stock |
|  | SS63 | Lab stock |
|  | SS64 | Lab stock |
| *Pasteurella multocida* (*P. multocida*) | PM002 | Lab stock |
|  | PM008 | Lab stock |
| *Actinobacillus pleuropneumoniae* (*A. pleuropneumoniae*) | APP015 | Lab stock |
|  | APP016 | Lab stock |
|  | APP017 | Lab stock |
|  | APP018 | Lab stock |

ATCC, American Type Culture Collection

**Supplementary Table 5. The primer sequences used for the RT-qPCR.**

| Genes | Primer sequence (5’ to 3’) |
| --- | --- |
| 16S rDNA | Forward: AGAGTTTGATCCTGGCTCAG |
|  | Reverse: GGTTACCTTGTTACGACTT |
| gyrB | Forward: ATGGCTATGGAACAGCAGCAAAATAG |
|  | Forward:AATATCAAGATTTTTCACGTATCTGGCGT |
| TNF-α | Forward: CCACGCTCTTCTGTCTACTG |
|  | Reverse: ACTTGGTGGTTTGCTACGA |
| IL-1β | Forward: ACCTGTGTCTTTCCCGTGG |
|  | Reverse: TCATCTCGGAGCCTGTAGTG |
| IL-6 | Forward: GAGCCCACCAAGAACGATA |
|  | Reverse: TTGTCACCAGCATCAGTCC |
| IL-10 | Forward: TGGACAACATACTGCTAACCG |
|  | Reverse: GGGCATCACTTCTACCAGGT |
| ZO-1 | Forward: CTGGTGAAGTCTCGGAAAAATG |
|  | Reverse: CATCTCTTGCTGCCAAACTATC |
| Occludin | Forward: CAGGATGCCAATTACCATCAAG |
|  | Reverse: GGGTTCACTCCCATTATGTACA |
| Claudin | Forward: AGATACAGTGCAAAGTCTTCGA |
|  | Reverse: CAGGATGCCAATTACCATCAAG |
| MUC2 | Forward: CGAGCACATCACCTACCACATCATC |
|  | Reverse: TCCAGAATCCAGCCAGCCAGTC |
| β-actin | Forward: GACCTCTATGCCAACACAGT |
|  | Reverse: CACCAATCCACACAGAGTAC |
| GAPDH | Forward: TGTTCCTACCCCCAATGTGT |
|  | Reverse: GGTCCTCAGTGTAGCCCAAG |

**Supplementary Table 6. Disease activity index (DAI) parameters and their associated scoring schemes.**

| **Score** | **Weight loss (%)** | **Stool consistency** | **Blood in stool** |
| --- | --- | --- | --- |
| 0 | None | Normal | Normal |
| 1 | 1-5 | Slightly loose stool | Small presence of blood |
| 2 | 5-10 | Loose stool | Significant presence of blood |
| 3 | 10-15 | Diarrhea | Gross blood |
| 4 | >15 |  |  |
